# Structure and Stability of the designer protein WRAP-T and its permutants

**DOI:** 10.1101/2021.04.09.438948

**Authors:** Bram Mylemans, Xiao Yin Lee, Ina Laier, Christine Helsen, Arnout R.D. Voet

## Abstract

*β*-Propeller proteins are common natural disc-like pseudo-symmetric proteins that contain multiple repeats (‘blades’) each consisting of a 4-stranded anti-parallel *β*-sheet. So far, 4- to 12-bladed *β*-propellers have been discovered in nature showing large functional and sequential variation. Using computational design approaches, we created perfectly symmetric *β*-propellers out of natural pseudo-symmetric templates. These proteins are useful tools to study protein evolution of this very diverse fold. While the 7-bladed architecture is the most common, no symmetric 7-bladed monomer has been created and characterized so far. Here we describe such a engineered protein, based on a highly symmetric natural template, and test the effects of circular permutation on its stability. Geometrical analysis of this protein and other artificial symmetrical proteins reveals no systematic constraint that could be used to help in engineering of this fold, and suggests sequence constraints unique to each *β*-propeller sub-family.

## 1 Introduction

*β*-propeller proteins can be found in all domains of life. They consist of repeating units adopting a circular shape around a central channel. These repeat units or “blades” consist of four-stranded anti-parallel *β*-sheets. The number of repeats varies between four to twelve, with seven being the most common [Murzin, 1992]. Hydrophobic interactions contribute the most to the stability of the fold, but in addition, most propellers possess a closing mechanism. In the four-bladed variants there is often a disulfide bridge between the N-terminal and C-terminal blade. In the larger propellers a so-called “Velcro closure” is found, in which the N- and C-terminal *β*-strands are part of the same *β*-sheet, keeping the propeller closed. This “Velcro” can occur in different positions; in the “1+3 Velcro” the closing *β*-sheet consists of one N-terminal and three C-terminal strands, while for the “3+1 Velcro” the opposite is true. Both of these configurations are common in nature [Fülöp and Jones, 1999, Chen et al., 2011a].

Like other repeat proteins, *β*-propellers are pseudo-symmetrical, with similar but not identical repeated motifs, suggesting evolution through multiple cycles of duplication and subsequent divergence. It is therefore believed that repeat proteins have evolved from smaller ancestral units, with diversification of the repeating units leading to the present natural sequences [Chaudhuri et al., 2008, Kopec and Lupas, 2013]. This process appears to have occurred independently many times, resulting in the existence of different distinct repeating motifs within the *β*-propeller family. The most archetypical are the WD40 repeats, but others including the Asp-box and Kelch repeats are also widespread [Chaudhuri et al., 2008]. While these families generally prefer a specific number of repeats, variations do occur. WD40 proteins for example are typically seven-fold symmetric, but six- and eight-fold proteins have also been observed [Hu et al., 2017].

Repeat proteins are a favoured target for protein design because larger domains can be constructed from a smaller subset thus reducing the design space considerably [Main et al., 2005]. Many of these studies aim to increase our understanding of protein evolution [Mylemans et al., 2021]. Pizza was the first successfully designed symmetrical *β*-propeller by reverse engineering the duplication-fusion process. Ancestral repeat sequences calculated from a natural protein were computationally mapped onto a designed symmetric backbone. This resulted in a symmetric *β*-propeller comprised of six identical blades [Voet et al., 2014]. Interestingly the protein can self-assemble from smaller fragments, validating these as probable evolutionary intermediates. The self-assembling properties of Pizza were further explored by engineering a metal-binding site into the Pizza protein, allowing it to scaffold a cadmium chloride nanocrystal, opening up possibilities for symmetric *β*-propeller proteins in the field of biomineralization.[Voet et al., 2015, Shang and Nienhaus, 2015] Other Pizza derivatives exhibited catalytic activity [Clarke et al., 2019], bound inorganic metaloxoclusters [Vandebroek et al., 2020], or were joined with *α*-helices to create protein cages [Vrancken et al., 2020]. As symmetric proteins are highly interesting for the creation of symmetric macromolecular assemblies, the same computational design strategy was subsequently used to make the eight-bladed propeller, Tako8 [Noguchi et al., 2019] and the nine-bladed propeller Cake9 [Mylemans et al., 2020a]. The latter can also fold as an eight-bladed propeller when only eight repeats are expressed. This structural plasticity may explain the origin of abundantly present odd-numbered *β*-propeller through duplication and fusion events, but it limits the applicability of Cake9 as a building block for larger assemblies. An attempt was made to create a protein that can only adopt the nine-bladed propeller fold by designing a three-fold symmetric protein instead of a nine-fold symmetric one. This resulted in the Scone protein which interestingly folds as a permuted eight-bladed propeller despite nine repeats are expressed in the open reading frame [Mylemans et al., 2021]. These unexpected observations highlight the need for further optimization of the computational design strategies in order to accurately control the folding symmetry of *β*-propellers.

In this study, we aimed to investigate whether there are geometric patterns linking the 3D structure to the fold symmetry. Perfectly symmetric *β*-propellers ease this analysis compared to natural pseudo-symmetric propellers. With Pizza6, Tako8, Cake8 and Cake9 we have six-, eight-, and nine-bladed proteins at hand. A five-bladed symmetric propeller was designed by the group of Tawfik. They selected functional proteins from an ancestral sequence library constructed from the sugar binding protein Tachylectin-2, resulting in a sequence of 47 amino acids that multimerized into a functional five-bladed propeller [Smock et al., 2016]. The group of Lupas more recently reported a highly repetitive propeller with 14 repeats folding as two seven-bladed propeller domains (PDB:2ymu) in *Nostoc punctiforme*. They named this protein “WD40-family Recently Amplified Propeller (WRAP)”. Similar to Cake however this sequence possesses the ability to adopt both multimeric eight- and nine-fold symmetry, when fragments are expressed [Afanasieva et al., 2019].

However, in nature the most commonly observed number of blades is seven, and so far no monomeric 7-bladed symmetrical propeller protein has been reported. Here, we report the crystallographic structure of a seven-fold repeating monomeric protein, based on 2ymu. We also sought to investigate the influence of the “Velcro” positioning on the stability of the protein [Mylemans et al., 2020b]. As a full set of designed symmetric *β*-propellers from five to nine-fold symmetry is now available, we also analysed and compared their geometry in order deduce rules which may be used as guidelines or restraints in future designs.

## 2 Results

We began our project before a consensus sequence of the 14 WRAP blades was reported by the group of Lupas Afanasieva et al. [2019]. Therefore we independently aligned all repeats and identified the most common amino acid at each position. The WRAP repeats show almost perfect conservation with very few mutations, and the only position which is not highly conserved is the ninth residue of the inner *β*-strand, where arginine and tryptophan occur equally often. As neither amino acid dominated, we did not choose either. Choosing an arginine may have yielded a supercharged structure at the core potentially preventing stable folding, while choosing the tryptophan could have depleted the supply of this single codon amino acid during expression and therefore negatively influence the yield of the protein. Hence, we constructed a sequence logo based on 7,142 sequences classified in the PROSITE database as WD40 repeats [Sigrist et al., 2012]. In agreement with previous studies [Jain and Pandey, 2018, Xu and Min, 2011], we observed threonine to be the most commonly observed residue at this position. As this manually edited consensus sequence is only different in one residue compared to the WRAP structures later reported by the Lupas group [Afanasieva et al., 2019], we refer to the here reported sequence as WRAP-T. The template protein, repeat alignment and sequence logo are shown in figure 1.

**Figure 1:**
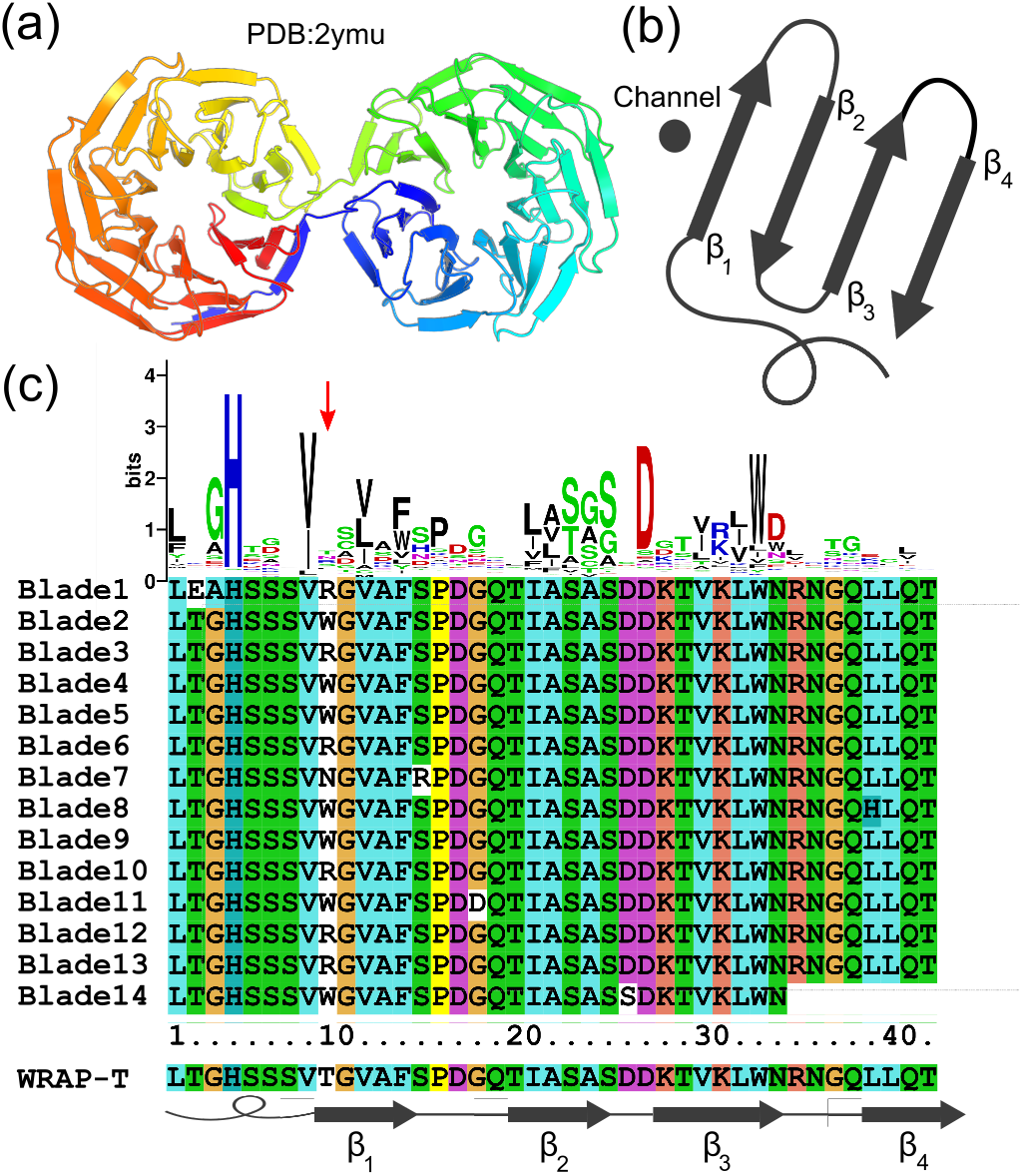
Design of the WRAP-T repeat. The structure 2ymu, the protein consists of 14 nearly identical WD40 repeats, folded into two seven-bladed *β*-propellers (a). A schematic representation of a single WD40 repeat. The repeat folds in four *β*-strands numbered form one to four from the inside of the cavity to the outside of the cavity (b). Sequence alignment of all 14 repeats of 2ymu, the sequence logo on top was constructed based on 7,142 sequences classified in the PROSITE database as WD40 repeats. The sequence of WRAP-T is shown on the bottom accompanied by a diagram showing the location of the *β*-strands (c).

In order to investigate the influence of the “Velcro closure” position we designed four synthetic DNA fragments expressing circularly permuted versions of the protein : nvWRAP-T without a “Velcro strap”, v31WRAP-T with a “Velcro” close to the central cavity, v22WRAP-T with a “Velcro” in the middle of the four *β*-strands and v31WRAP-T with the outer “Velcro closure” (also present in the model protein 2ymu). A model and diagram of the first repeat are shown in figure 2.

**Figure 2:**
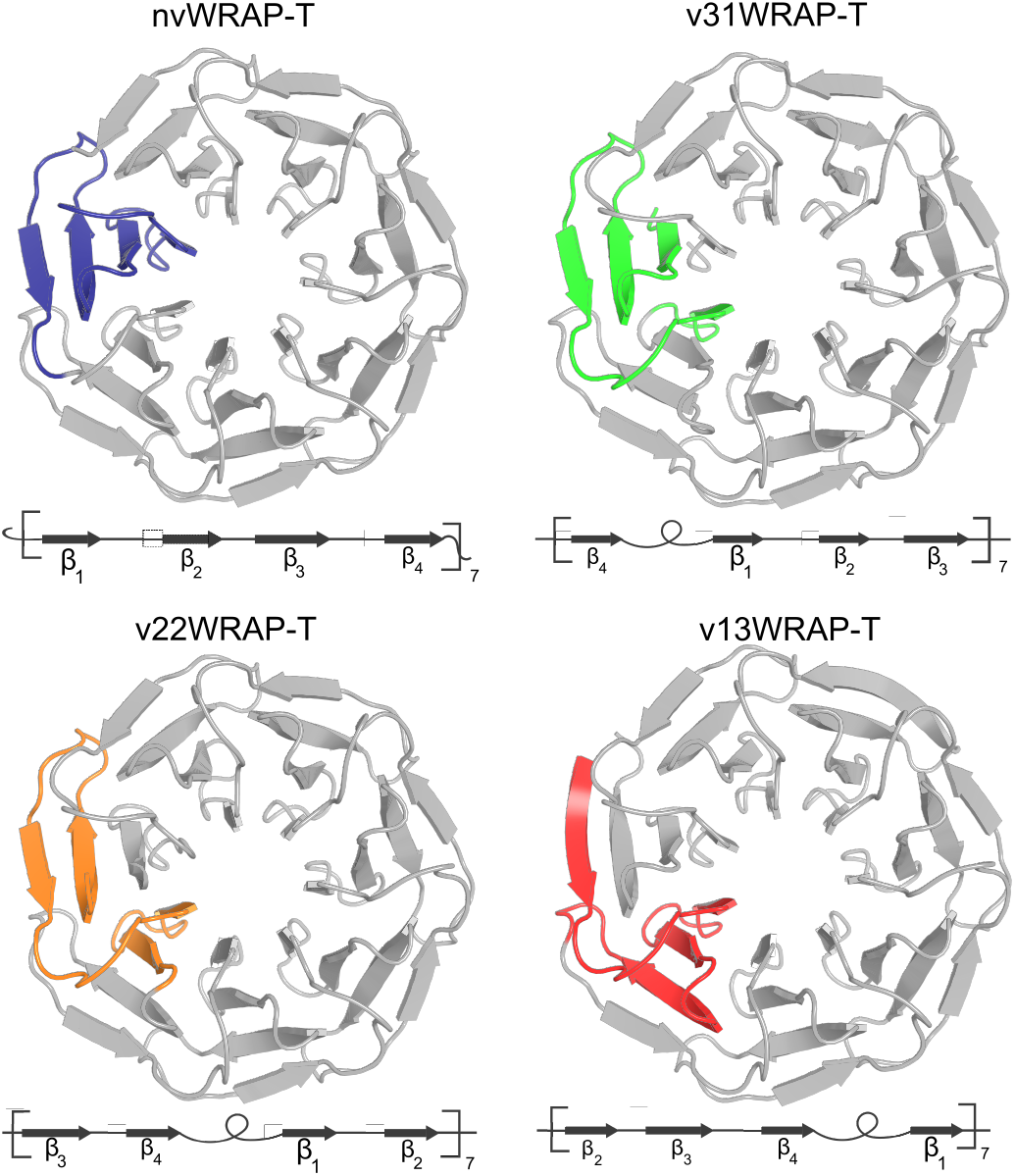
Construction of the “Velcro” variants, each “Velcro” protein has the same sequence but the N-terminal amino acid is shifted within the repeat thus resulting in a different location of the “Velcro”. The first repeat is shown in colour, underneath each structure a diagram is shown illustrating the order of *β*-strands within the repeat.

The DNA fragments (table S1) were cloned into a pET28 vector, and expressed in *E. coli*. Purification with a Ni-NTA column and size-exclusion chromatography yielded protein of high purity, suitable for crystallization. Each protein variant was monodisperse and eluted at the same volume, indicating the same molecular size, which was verified using analytical gelfiltration, see figure 3. Optimal crystallization conditions were determined using standard screens. Each protein crystallized in a different condition (table S2). High quality X-ray diffraction data-sets (see table 1) were obtained with high resolution limits between 1.4 and 1.8 Å. Initial models were obtained using molecular replacement, starting with a propeller domain from PDB entry 2ymu. While the space group and packing differ between the crystals (figure S1), the structure is identical for each protein. The backbone RMSD between each artificial protein and the 2ymu template is below 0.4 Å. The crystal structures also show that the proteins are highly internally symmetric with a backbone RMSD between individual repeats below 0.3 Å. The characteristic hydrogen bridge network in WD40 repeats, between a conserved histidine, serine, aspartate and tryptophan is clearly visible in the electron density, see figure 4. In order to investigate the influence of the “Velcro” position on stability, the CD signal at 218 nm was monitored while raising the protein temperature, see figure 3. The resulting curves were fitted with a Boltzmann-sigmoid to determine the melting temperature and ΔH, see table 2. Thermal reversibility was tested by heating the samples to 85°C and then back to 20°C to compare the spectrum (figure S2). Chemical stability was tested by measuring intensity changes in trypthophan fluorescence at the 320 nm wavelength under increasing concentrations of guanidine hydrochloride (GdnHCl). From this, the fraction of denatured protein could be calculated by assuming the protein is completely unfolded at the highest concentration. When possible this data was fitted with a Boltzmann-sigmoid curve to determine the concentration at which 50% of the protein is denatured and the proportionality constant (m), see table 2. v13WRAP-T remained stable at high GdnHCl concentration thus fitting the data was not possible.

**Figure 3:**
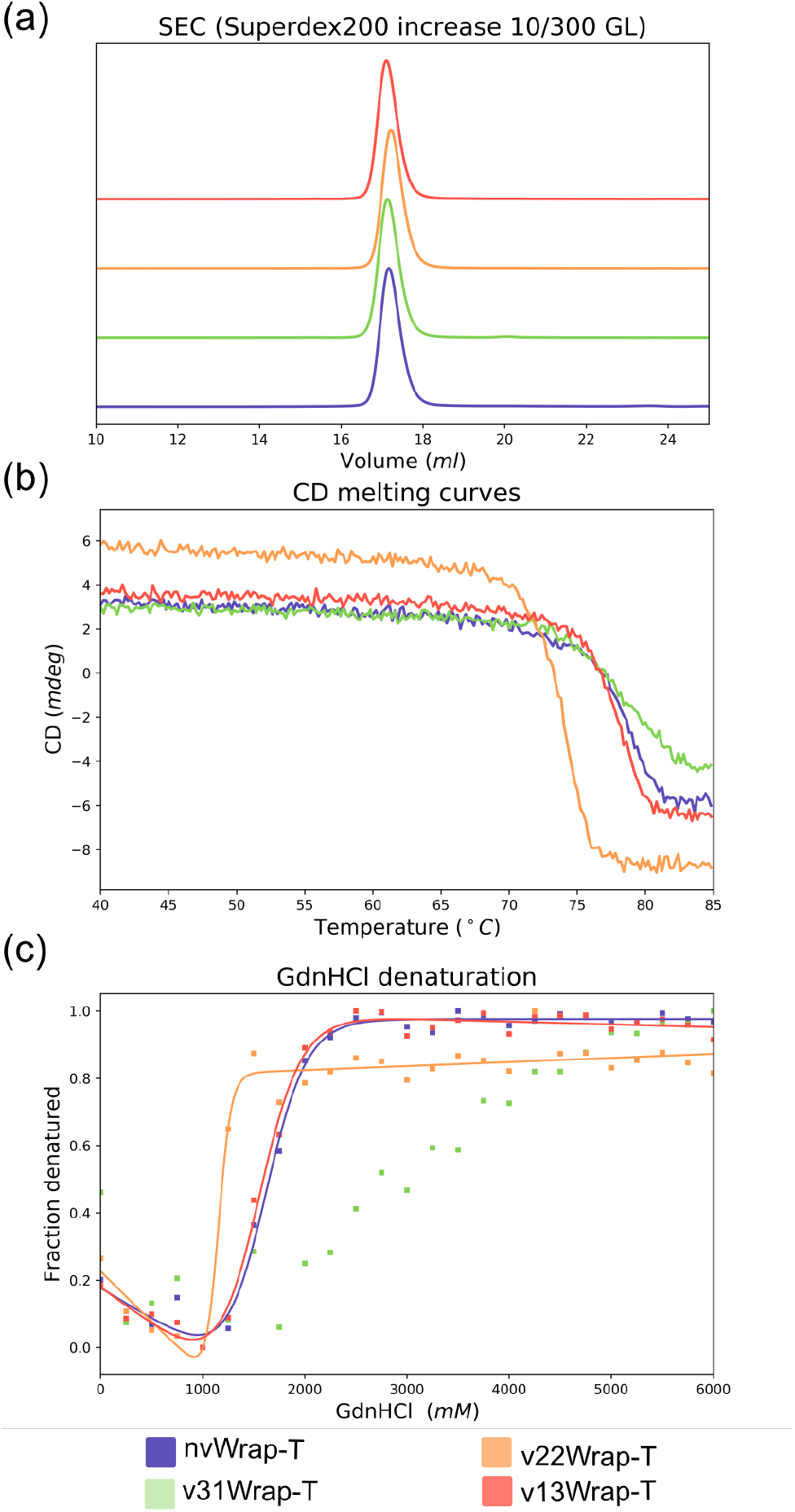
Purification and stability data for the WRAP-T proteins. Normalized analytical size exclusion indicates all proteins have the same hydrodynamic volume (a). Measuring the CD signal at 218 nm while increasing the temperature shows the melting point of each protein see data table 1 (b). The fluorescent signal of tryptophan was followed at 320 nm this allows for the calculation of fraction denatured under increasing Guanidine hydrochloride concentration (c).

**Table 1:** Data collection and refinement statistics of X-ray structures. Values in parentheses are for the outer shell.

|  | nvWRAP-T | v31WRAP-T | v22WRAP-T | v13WRAP-T |
| --- | --- | --- | --- | --- |
| **PDB entry** | **7BIE** | **7BID** | **7BIF** | **7BIG** |
| **Data collection** |  |  |  |  |
| Diffraction source | I03, DLS | I04, DLS | I03, DLS | I03, DLS |
| Wavelength (Å) | 0.9763 | 0.9763 | 0.9763 | 0.9763 |
| Resolution range (Å) | 59.76 - 1.80 (1.84 - 1.80) | 44.64 - 1.80 (1.84 - 1.80) | 29.47 - 1.40 (1.42 - 1.40) | 52.13 - 1.8 (1.84 - 1.8) |
| Space group | $C2$ | $P2_1P2_1P2_1$ | $P2_1$ | $P2_1P2_1P2_1$ |
| $a, b, c$ (Å) | 79.66, 48.497, 121.95 | 44.64, 61.40, 90.41 | 63.88, 72.89, 93.6 | 64.93, 81.48, 87.42 |
| $\alpha, \beta, \gamma$ (°) | 90, 101.47, 90 | 90, 90, 90 | 90, 96.01, 90 | 90, 90, 90 |
| Reflections (measured/unique) | 276, 861/41, 838 | 222, 824/26, 008 | 1, 073, 302/166, 377 | 567, 324/43, 750 |
| Completeness (%) | 98.6 (97.5) | 99.8 (97.3) | 99.2 (86.5) | 100 (100) |
| Mean $I/\sigma(I)$ | 11.9 (5.5) | 55.4 (21.6) | 25.5 (6.2) | 15.4 (2.5) |
| Multiplicity | 6.6 (6.2) | 8.6 (5.0) | 6.5 (4.9) | 13.0 (12.2) |
| $R_{\text{pim}}$ | 0.039 (0.114) | 0.009 (0.019) | 0.016 (0.094) | 0.028 (0.281) |
| CC(1/2) | 0.993 (0.959) | 1.000 (0.998) | 1.000 (0.969) | 0.997 (0.915) |
| Wilson B factor (Å<sup>2</sup>) | 16.7 | 11.32 | 8.8 | 20.2 |
| **Refinement Statistics** |  |  |  |  |
| $R$ factor/$R_{\text{free}}$ | 0.173/0.216 | 0.1476/0.1827 | 0.156/0.181 | 0.195/0.243 |
| No. of atoms in structure |  |  |  |  |
| Protein | 4, 153 | 2, 153 | 8, 461 | 4, 129 |
| Ligand | 13 | 0 | 0 | 0 |
| Water | 513 | 326 | 1361 | 391 |
| R.m.s. deviations from ideal |  |  |  |  |
| Bond lengths (Å) | 0.007 | 0.016 | 0.006 | 0.004 |
| Bond angles (°) | 0.995 | 1.34 | 1.010 | 0.727 |
| Chiral volumes (Å<sup>3</sup>) | 0.069 | 0.101 | 0.101 | 0.054 |
| Ramachandran plot, residues in (%) |  |  |  |  |
| Most favorable region | 95.25 | 95.40 | 94.98 | 95.77 |
| Additional allowed region | 4.75 | 4.58 | 5.02 | 4.23 |
| Average $B$ factor (Å<sup>2</sup>) | 18.0 | 14.0 | 13.0 | 27.0 |

**Figure 4:**
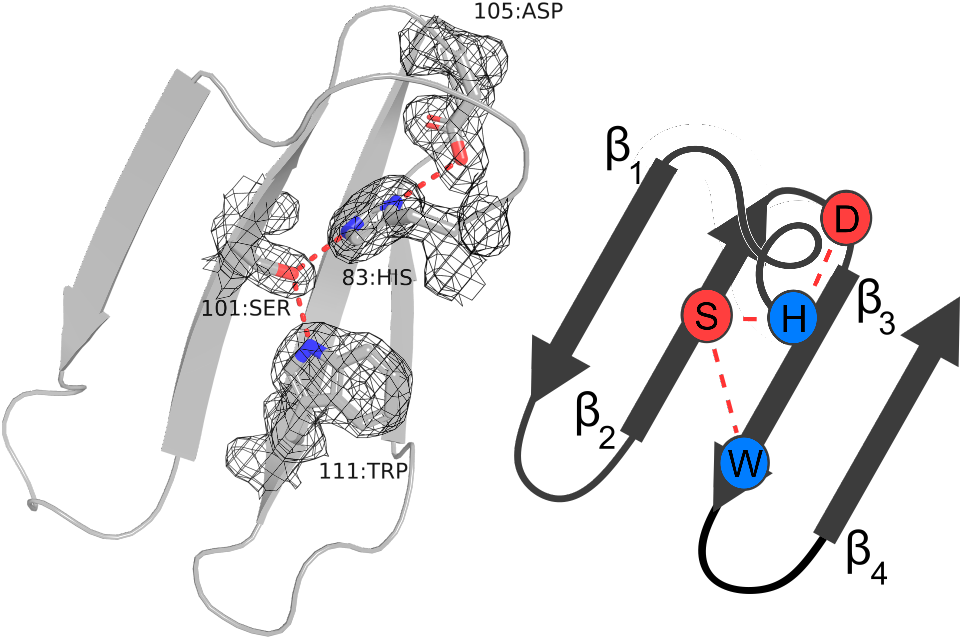
The hydrogen bonding network in WD40 repeats. A single repeat from the crystal structure nvWRAP-T showing the highly conserved hydrogen bonding network between 111:Trp, 101:Ser, 83:His and 105:Asp. The electron density is shown as a *F*_*c*_ – *F*_*o*_ map at sigma level 1. Beside the structure a diagram indicating the hydrogen network is shown.

**Table 2:** Stability parameters of the proteins

|  | nvWRAP-T (CD) | v31WRAP-T | v22WRAP-T | v31WRAP-T |
| --- | --- | --- | --- | --- |
| $T_m$ (°C) | $78.33 \pm 0.06$ | $78.63 \pm 0.09$ | $73.80 \pm 0.03$ | $77.68 \pm 0.03$ |
| $\Delta H_m$ (kJ/mol) | $-590 \pm 17$ | $-430 \pm 11$ | $-860 \pm 16$ | $-790 \pm 16$ |
| $\Delta\Delta G_{D-N}$ (kJ/mol) | $-1.5 \pm 0.01$ | $-2.1 \pm 0.2$ | $8.7 \pm 0.2$ | / |
| $\Delta G_0^{H_2O}$ (kJ/mol) | $2.21 \pm 0.04$ | / | $1.17 \pm 0.03$ | $2.05 \pm 0.05$ |
| m value (kJ $\text{mol}^{-1} \text{M}^{-1}$) | $10 \pm 1.3$ | / | $28 \pm 6$ | $9 \pm 1.3$ |
| $C_m$ (M) | $2.21 \pm 0.04$ | / | $1.17 \pm 0.03$ | $2.05 \pm 0.05$ |
| $\Delta\Delta G_{D-N}$ (kJ/mol) | $-1.5 \pm 0.1$ | / | $16 \pm 4$ | / |

## 3 Discussion

A symmetric seven-bladed *β*-propeller was designed from a highly symmetric protein with unknown function (PDB:2ymu). This design completes the arsenal of designed symmetrical *β*-propellers from five- to nine-fold symmetry. All these proteins consist of a motif between 40 and 49 amino acids long that is repeated multiple times. Some motifs permit multiple symmetries. The Cake sequence for example can both fold into an eight- and nine-bladed propeller depending on the number of repeats expressed [Mylemans et al., 2020a], and the WRAP sequence has also been shown to possess this property [Afanasieva et al., 2019]. Most sequences however will only properly fold in a specific symmetry; Pizza will only adopt a six-bladed propeller, even when seven repeats are tandemly placed in a single polypeptide Voet et al. [2014]. So far it has proved impossible to predict accurately the correct folding symmetry for a designed sequence, or to assess whether a designed structure will correctly fold. In a recent study we designed a nine-bladed propeller that was found in practice to form an eight-bladed propeller. In order to design confidently a *β*-propeller *ab initio*, improved understanding of these structures is required. Analysis of artificial as well as natural proteins will assist in this process.

All the WRAP-T permutants expressed well and could be purified easily. Solving the crystal structures revealed that they possess an identical structure. Interestingly, the different position of the chain termini had an influence on the preferred crystallisation conditions and crystal packing. While for previously designed symmetric proteins, the proteins typically packed in distinct symmetric ways reflecting the inherent six-, eight- or nine-fold symmetry of the proteins, here no clear symmetric interactions could be observed within the lattice. This may be expected given the incompatibility of sevenfold internal symmetry with any space group. Despite the non-equivalent positions of each blade within the crystal structures, the proteins proved to be perfectly symmetric, with a TM-score above 0.97 calculated by CE-symm [Bliven et al., 2019]. Analysing the thermal and chemical stability of the WRAP-T permutants revealed that nvWRAP-T, lacking the velcro closure, is just as stable as the variant v13WRAP-T. While v31WRAP-T is considerably more resistant to GdnHCl induced denaturation compared to the other proteins, its increased thermal stability is not that significant. As expected, v22WRAP-T is the least stable protein, reflecting the fact this conformation is almost never seen in nature.

In a previous study we investigated the influence of the “Velcro” strap on the stability of the Pizza protein, which belongs to the NHL family of propeller proteins [Mylemans et al., 2020b]. It was found that the two“non-Velcro” Pizza variants (respectively having the termini just in front of the first strand, or behind the last strand of the blade) are the least stable, having a melting temperature of 41.78°*C* and 45.32°*C* compared to 52.48°*C* for the regular Pizza. The Velcro2-2 variant is the least stable Pizza with a “Velcro”, having a melting temperature of 44.13°*C*. The Velcro3-1 variant is almost as stable as the regular Pizza, with a melting temperature of 52.48°*C*. The same trends can be found for chemical denaturation, with nv1Pizza6 having a *C*_*m*_ value of 0.660 *M*, nv2Pizza6 a value of 0.782*M* and v22Pizza6 a value of 0.749 *M*, this can be compared to a value of 0.949 *M* for the regular Pizza. Just as with WRAP-T, the inside “Velcro3-1” position is highly resistant to chemical denaturation.

In both Pizza and WRAP-T, the 2-2 “Velcro” is the least stabilising, which presumably explains why it is rarely observed in nature. The few exceptions include the RCC1-like domains Renault et al. [1998], which are only distantly related to other *β*-propellers Chaudhuri et al. [2008]. For WD40 repeats this decreased stability can be explained through the disruption of the highly conserved hydrogen bonding network between a serine/threonine in *β*-strand two, an aspartate and tryptophan in *β*-strand three and a histidine in the connecting loop, see figure 4. The importance of this conserved hydrogen bonding network for the WD40 class of proteins was previously confirmed by mutating the amino acids involved Wu et al. [2010]. However the Pizza protein does not posses this network, yet the middle “Velcro” is still the least stable. Possibly burying the charged chain termini deep within the protein is destabilising enough to explain the observed results. In both cases the “Velcro3-1” position yields the protein with the highest stability, especially against chemical denaturants such as GdnHCl. A striking difference however is the stability of the “non-Velcro” variant, the least stable for Pizza but as stable as the natural “Velcro” position in WRAP-T. This may be due to differences between the sequences, as the two different “non-Velcro” variants of Pizza have different stabilities as well. However, these two examples alone may not be sufficient to determine a clear correlation, and additional proteins will be needed to confirm whether the inner “Velcro” is consistently the most stable conformation for propeller proteins. Such information may be helpful in engineering *β*-propeller enzymes for industrial applications such as the production of fructans by fructosyltransferases [Flores-Maltos et al., 2016].

Another goal of this research was the investigation of common trends in the backbone arrangement, especially the blade orientation within the propeller architecture. These findings could serve as guidelines for the *de novo* design of propeller backbones with a desired symmetry. We performed a structural analysis of the currently reported perfectly symmetric monomeric propeller proteins, but excluded any non-symmetric proteins in order to avoid bias to the backbone structure induced by the pseudo-symmetry of the sequence. The investigated proteins include: Tachylectin-2 (PDB:5c2m), Pizza (PDB:3ww9), WRAP-T (PDB:7big), Tako8 (PDB:6g6n), Cake8 (PDB:6tjg), and Cake9 (PDB:6tjh). Each has a different number of repeats, with the Cake sequence giving both eight- and nine-fold symmetric variants. A structural alignment of a single repeat of each protein allowed for the selection of four reference *C*_*α*_ atoms in each repeat marked in the structure and sequence alignment in figure 5. Two are located on the inside *β*_1_-strand, the other two are found in the *β*_3_-strand. From these reference points we calculated six distances and four angles to characterize the specific protein. The distances from the inner position to the central axis (a,b); the distance between reference points in subsequent repeats, for inner points (c,d) and outer points (e,f); the rotation angle (*α*); the twist angles of the blade as the dihedral angle between the *β*_1_-strand and the central axis (*β*) and the internal dihedral angle between the two strands (*γ*); and finally the tilt angle between the central axis and the *β*_1_-strand (*δ*). Average values and standard deviation of these values are shown in table 3, and plots are shown in figure 5. Both WRAP-T and Tako belong to the WD40 family of proteins and are marked by a hollow triangle, while the other propellers belong to different families and are marked by a circle. As could be expected, the rotation angle *α* is entirely dictated by the symmetry 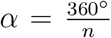, with *n* the number of blades. The channel radius is also highly dependent on symmetry, generally increasing with increasing blade number. Tachylectin-2 is the exception, having a larger central channel than Pizza. A linear regression fit of the channel radii against symmetry number indicates a high correlation (*r*^2^ > 0.95) for the larger propellers, but only a weak correlation if Tachylectin-2 is included.

**Table 3:** Geometric parameters of symmetric designer proteins

|  | Tachylectin-2 | Pizza6 | WRAP-T | Tako8 | Cake8 | Cake9 |
| --- | --- | --- | --- | --- | --- | --- |
| a (Å) | $7.0 \pm 1.0$ | $4.6 \pm 0.2$ | $5.82 \pm 0.06$ | $7.32 \pm 0.06$ | $6.63 \pm 0.06$ | $8.34 \pm 0.04$ |
| b (Å) | $6.28 \pm 0.04$ | $7.1 \pm 0.3$ | $8.51 \pm 0.14$ | $11.4 \pm 0.7$ | $10.85 \pm 0.05$ | $12.96 \pm 0.11$ |
| c (Å) | $8.19 \pm 0.04$ | $4.5 \pm 0.4$ | $5.03 \pm 0.11$ | $5.6 \pm 0.2$ | $5.08 \pm 0.05$ | $5.74 \pm 0.07$ |
| d (Å) | $7.41 \pm 0.04$ | $7.2 \pm 0.2$ | $7.38 \pm 0.14$ | $8.74 \pm 0.06$ | $8.30 \pm 0.09$ | $8.9 \pm 0.2$ |
| e (Å) | $19.8 \pm 0.16$ | $14.7 \pm 0.3$ | $13.32 \pm 0.15$ | $13.2 \pm 0.13$ | $13.1 \pm 0.1$ | $12.92 \pm 0.03$ |
| f (Å) | $18.7 \pm 0.9$ | $16.24 \pm 0.10$ | $15.1 \pm 0.2$ | $15.0 \pm 0.3$ | $14.58 \pm 0.12$ | $14.43 \pm 0.08$ |
| $\alpha$ (°) | $70 \pm 7$ | $60 \pm 5$ | $51.4 \pm 1.3$ | $45.0 \pm 1.8$ | $45 \pm 0.4$ | $40.0 \pm 0.5$ |
| $\beta$ (°) | $-10 \pm 6$ | $-4.8 \pm 1.3$ | $-0.7 \pm 0.3$ | $-9 \pm 3$ | $11.1 \pm 0.3$ | $14.0 \pm 0.4$ |
| $\gamma$ (°) | $-22.84 \pm 0.18$ | $-29 \pm 2$ | $-32.8 \pm 0.5$ | $-23.8 \pm 1.5$ | $-44.4 \pm 0.4$ | $-45.9 \pm 0.7$ |
| $\delta$ (°) | $11 \pm 6$ | $14.8 \pm 1.1$ | $14.5 \pm 0.8$ | $25 \pm 4$ | $28.2 \pm 0.6$ | $13.0 \pm 0.1$ |

**Figure 5:**
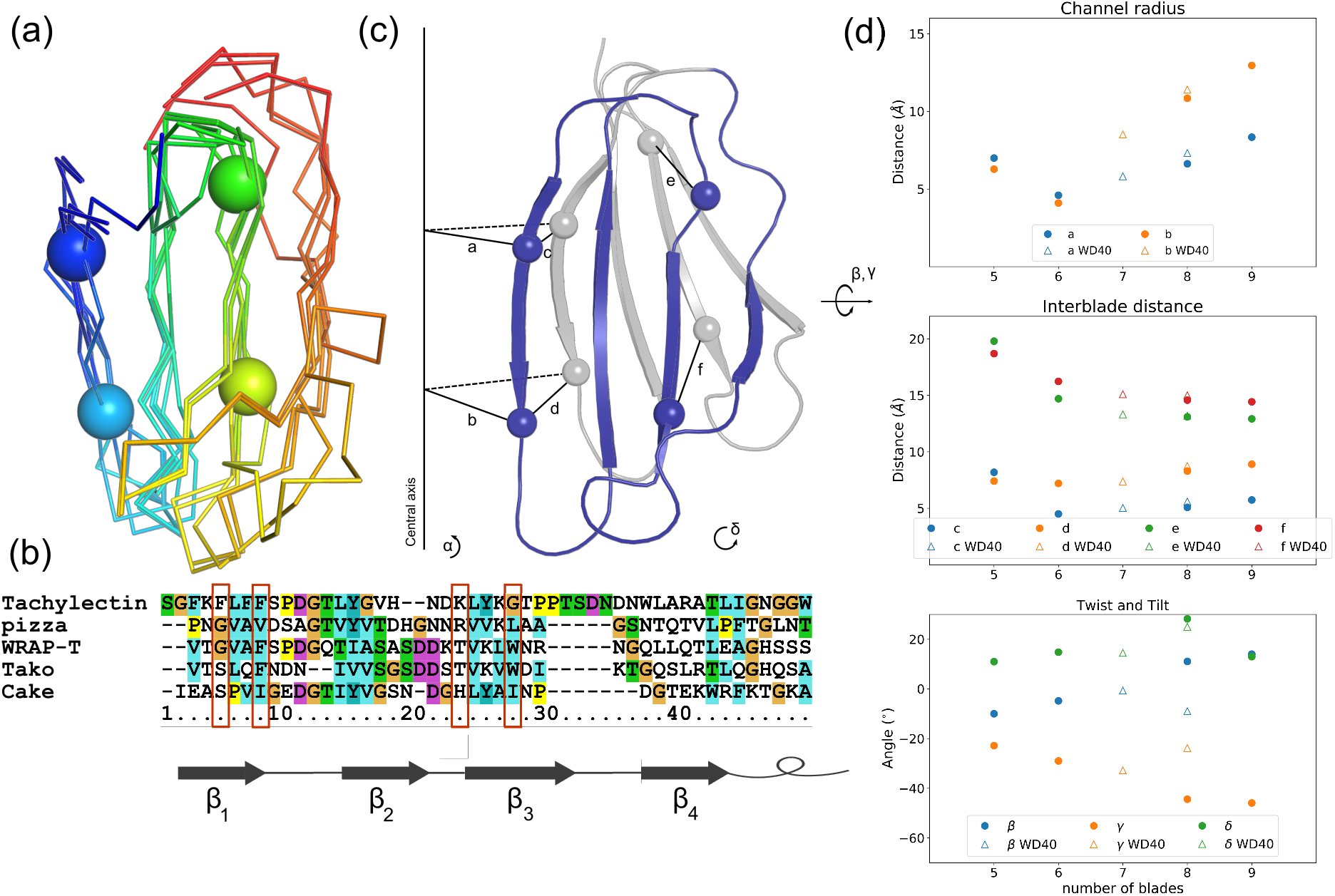
Geometric analysis of different symmetric designer proteins: The five-fold Tachylectin-2 (PDB:5c2m), the six-fold Pizza (PDB:3ww9), seven-fold WRAP-T (PDB:7big), eight-fold Tako (PDB:6g6n) and Cake (PDB:6tjg) and the nine-fold Cake (PDB:6tjh). An alignment of a single blade from each protein. The spheres indicate the location of the chosen reference point (a). A structure based sequence alignment of a single blade from each protein, the red boxes show the location of the sphere reference point in each sequence (b). A cartoon of two subsequent blades with spheres indicating the reference points. The chosen parameters are shown: The distances from the inner position to the central axis (a,b); The distance between reference points in subsequent repeats, for inner points (c,d) and outer points (e,f); The rotation angle (*α*); The twist angles of the blade as the dihedral angle between the *β*_1_-strand and the central axis (*β*) and the internal dihedral angle between the two strands (*γ*); And the tilt angle between the central axis and the *β*_1_-strand (*δ*) (c). Graphs showing the average value for each parameter in function of repeat number. WRAP-T and Tako are both WD40 repeats and are represented by a hollow triangle, the other proteins belong to different families and are shown as a solid circle (d).

The inter-blade distances also follow a linear pattern; while the inside inter-blade distance increases with the number of blades, the outside distance decreases. Again Tachylectin is the one exception to this rule. It should be noted that these distances are not independent, and using the simple model of an isosceles triangle, 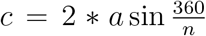, the same is true for the relation between d and b. A similar formula can be applied to e and f with the difference that the distance between the first and third *β*-strand has to be taken in to account. As can be seen from the alignment of all blades in figure 5, this distance is nearly identical for all symmetries, thus e and f are also completely determined by a/b and the number of repeats. This results in a linear decrease with *r*^2^ > 0.8 From these distances it is clear that Tachylectin-2 is the misfit within these proteins, although it is unclear whether this is true for all five-bladed propellers, or specific to Tachylectin-2 and its homologs. The dihedral twist angles show some relation with symmetry, with *β* increasing while the internal *γ* angle decreases, however *r*^2^ is only above 0.5. In this case Tako seems to be the misfit, and excluding this protein yields a much higher correlation, *r*^2^ > 0.95. The tilt angle *δ* seems uncorrelated to the blade number with an *r*^2^ < 0.2. It is interesting to note that for the two Cake proteins, the dihedral angles are nearly identical while the tilt angles are different. It seems that this sequence can adopt different symmetry by increasing the channel radius and decreasing the tilt with respect to the central axis, similar as how a wire helix can be expanded or contracted in a similar way by rotating coils at each end in opposite directions about the central axis.

While the inter-blade distances obey simple geometric rules depending on the channel radius and symmetry, no design guidelines on blade orientation could be deduced from these few proteins. Tachylectin-2 shows that it is possible to have a larger channel with a low symmetry by including a longer loop between the third and fourth *β*-strand, filling the gap that would otherwise be created on the outside of the propeller. The Tako protein deviates from the trend and has drastically different angles from Cake8, the other eight-fold protein, showing that the same symmetry can be achieved by very different blade orientations.

It is known that close homologs of ring-shaped protein oligomers may have different numbers of subunits in the complex. For example the Tryptophan RNA-binding attenuation protein (TRAP) usually consists of eleven subunits, but the variant from *B. halodurans* (PDB: 3zzl) has twelve. This is caused by a deletion of the last five residues of each subunit, shifting the angle of rotation slightly. Equally the insertion of a larger amino acid on the inside of the circle also causes a shift in rotation angle, resulting in a conformation with twelve subunits [Chen et al., 2011b]. For the monomeric *β*-propellers, it seems more difficult to find such rules, as deviations in repeat structure are not pronounced and the same sequence can result in multiple symmetries.

## 4 Conclusion

Starting from a highly repetitive *β*-propeller sequence, we designed a consensus motif sequence that formed a perfectly symmetric 7-bladed *β*-propeller. This protein completes a set of artificial propeller proteins with five to nine-fold internal symmetry. A structural analysis of these proteins was performed to deduce their geometric patterns and rules. Although some trends are present in this small dataset, no definitive rules for propeller protein design can be deduced from it. A geometric relation between blade angle, radial distance and inter-blade distance was found, and a trend in angles of orientation could be observed, but exceptions were found to both tendencies. In the future it would be useful to compare proteins with the same symmetry but belonging to different propeller sub-families, in order to determine whether blade orientation is controlled more by the symmetry or features of the sequence.

## 5 Methods

### Protein preparation

Linear DNA sequences encoding the variants of WRAP-T were commercially prepared, amplified by PCR and inserted into pET28 vector using the NdeI and XhoI restriction sites. Proteins were expressed in *E. coli* BL21(DE3) and purified using published protocols [Noguchi et al., 2018]. Protein expression was induced by adding IPTG to a final concentration of 0.5 mM, and subsequently incubating the cells with shaking at 20°C for 20 h. Cells were harvested by centrifugation and suspended in 50 mM NaH_2_PO4, 200 mM NaCl and 10 mM imidazole. After lysis by sonication, cell debris was removed and the supernatant loaded onto a 10 ml volume nickel-sepharose column (GE Healthcare) equilibrated with the same buffer. The column was washed with a similar buffer containing 20 mM imidazole. The protein was finally eluted with buffer with 300 mM imidazole and digested with thrombin overnight at 4° C during dialysis into 50 mM NaH_2_PO4, 200 mM NaCl, 10 mM imidazole. The protease:WRAP-T ratio was 1:200. The protein was re-loaded onto the washed nickel-sepharose column and the same steps were repeated. The fractions containing tag-free WRAP-T were pooled before loading onto a Superdex-200 column (GE) equilibrated with 20 mM HEPES and 200 mM NaCl buffer at pH 8.0.

The purified proteins were concentrated to 10 mg/mL and shown to be at least 95% pure by SDS-PAGE. All purified proteins were analysed by size-exclusion chromatography (SEC). The SEC analysis was performed using a Superdex 200 increase 10/300 GL column (GH healthcare) equilibrated with 20 mM HEPES and 200 mM NaCl buffer at pH 8.0.

### Tryptophan fluorescence

Denaturation of the protein samples was measured by observing intrinsic tryptophan fluorescence with a Sapphire2 96-well plate reader (Tecan) following the protocol described in [Mylemans et al., 2020b]. 2 *μ*L of protein sample (OD_280_ of 10) were pipetted into 98 *μ*L of guanidinium hydrochloride (GdnHCl) solution to give a final protein concentration of 0.5 *μ*g/mL. GdnHCl concentration was tested from zero to 6 M, in steps of of 0.25 M. Tryptophan fluorescence was measured after seven days storage at 20°C, by observing the emission intensity at 330 nm after excitation at 280 nm. Samples were prepared and measured in triplicate [Moon and Fleming, 2011]. Fluorescence measurements from blank samples (containing no protein) were subtracted from the measured values. Normalized values for the fraction of denatured protein were obtained by scaling, with the maximum value set to 1 and the smallest to zero. The data points were fitted to a Boltzmann-sigmoid equation (1) with an in-house Python script utilising the *SciPy* library [Virtanen et al., 2020].

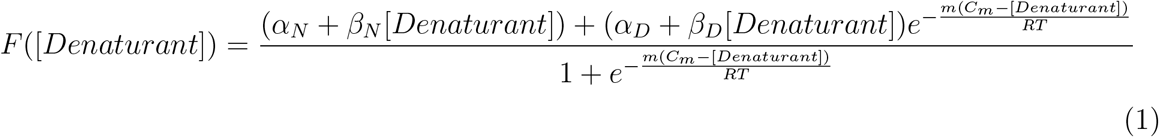

*F* is the measured fluorescence, *α*_*N*_ is the *F* value for the native state with zero denaturant, *β*_*N*_ is the slope of this signal, *α*_*D*_ and *β*_*D*_ are the corresponding values for the completely denatured state. These parameters are introduced because the denatured and native states change linearly with the denaturant concentration. *C*_*m*_ is the concentration at which 50% of the protein is denatured, m is the constant of proportionality 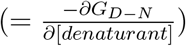 and has the dimensions of cal/mol/M [Clarke and Fersht, 1993].

The difference in free energy between the “Velcro” mutants and the natural position of v13WRAP-T was calculated with equation 2 Fersht [2017].

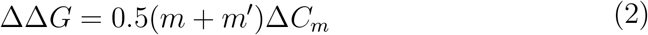

*m′* is the *m*-value of the mutant and Δ*C*_*m*_ is the difference in the concentration point at which 50% of the protein is denaturated.

### CD spectroscopy

All circular dichroism measurements were performed with a JASCO J-1500 instrument following the protocol described in [Mylemans et al., 2020b]. Thermal denaturation measurements were performed in a 2 mm path quartz cuvette using 0.05 mg/mL protein in 20 mM phosphate pH 7.6. The samples were heated from 20°C to 85°C in steps of 0.2°C while monitoring the CD signal at a wavelength of 218 nm.

The CD signal was fitted to a Boltzmann-sigmoid equation (3) using the same script employed for the chemical denaturation experiments.

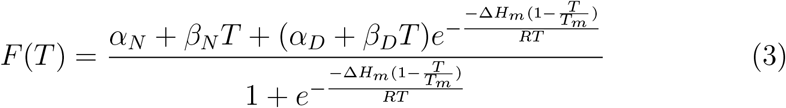

F is the measured signal (in this case the measured CD signal at 218 nm), *α*_*N*_ is the signal of the native state *α*_*D*_ and *β*_*D*_ are the corresponding values for the completely denatured state. *T*_*m*_ is the temperature at which 50% of the protein is denatured, and Δ*H*_*m*_ is the change of enthalpy upon denaturation [Becktel and Schellman, 1987]. The change in free energy between the “Velcro” mutants and the natural position of v13WRAP-T was calculated with equation 4 Fersht [2017].

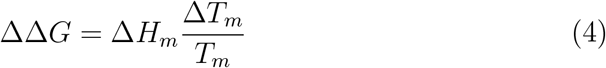

In which Δ*H*_*m*_ is the difference in enthalpy and Δ*T*_*m*_ the difference in melting temperature between the mutant and v13WRAP-T.

### Crystallisation and structure determination

The proteins were dialysed against 20 mM HEPES pH 8.0, and the concentration adjusted to 10.0 mg/mL. Crystallisation experiments were performed at 20°C using the sitting-drop vapor diffusion method with a Gryphon robot (Art Robbins). The crystals were washed in mother liquor containing glycerol, 20 % for nvWRAP-T, 25 % for v31WRAP-T, 15 % for v22WRAP-T and 10 % for v13WRAP-T as cryo-protectant before being flash-frozen and stored in liquid nitrogen. Data-sets were collected on the I03 or I04 beamlines at the Diamond Light Source (DLS) (Oxfordshire, UK). A total of 3600 images were taken with a 0.1° oscillation with a wavelength of 0.9763 Åusing an Eiger2 X 16M detector. The data were processed with XDS (for v31WRAP-T) [Kabsch, 2010a] or DIALS [Beilsten-Edmands et al., 2020], and scaling was carried out with Aimless [Kabsch, 2010b]. A single 7-bladed unit (Residues X to y) from parent template (PDB:2ymu) was used for molecular replacement with PHASER [McCoy et al., 2007]. Refinement was carried out with PHENIX.REFINE [Adams et al., 2010], with manual modifications performed with COOT [Emsley et al., 2010]. The CCP4 suite was used for general data handling [Collaborative et al., 1994]. An overview of data collection and refinement statistics is given in table 1. The structures are available from the PDB with entry codes 7bie, 7bid, 7bif, 7big for nvWRAP-T, v31WRAP-T, v22WRAP-T and v13WRAP-T, respectively.

## 6 Acknowledgements

We would like to thank Els Deridder for help with cloning. We thank the beamline staff at Diamond Light Source for their kind assistance. ARDV thanks Research Foundation Flanders for financial support (G0E4717N, G0F9316N and G051917N). BM thanks Research Foundation Flanders for a fellowship (GBM-D3229-ASP/17), ARDV and CH thank KU Leuven for C14/17/067 funding.

## Author contributions

BM designed the WRAP-T sequence, BM, XYL and IL purified and crystallised the proteins. XYL and BM analysed the diffraction data. BM performed the biophysical experiments. ARDV and CH conceived and supervised the experiments. All authors contributed to writing the manuscript

## Notes

### Competing Interest Statement

The authors have declared no competing interest.

